# E93 controls adult differentiation by repressing *broad* in *Drosophila*

**DOI:** 10.1101/2024.02.14.580315

**Authors:** Josefa Cruz, Enric Ureña, Xavier Franch-Marro, David Martín

**Affiliations:** Institute of Evolutionary Biology (IBE, CSIC-Universitat Pompeu Fabra), Passeig de la Barceloneta 37, 08003 Barcelona, Spain

**Author notes:** Institute of Healthy Ageing, Department of Genetics, Evolution and Environment, University College London, London WC1E 7JE, United Kingdom (Enric Ureña).

**Keywords:** Drosophila, wing development, E93, Broad, Chinmo, metamorphosis, enhancer regulation, temporal regulation, metamorphic transitions

## Abstract

In *Drosophila melanogaster*, successful development relies on the precise coordination of both spatial and temporal regulatory axes. The temporal axis governs stage-specific identity and developmental transitions through a number of genes, collectively forming the Metamorphic Gene Network. Among these, Ecdysone inducible protein 93F (E93) serves as the critical determinant for adult specification, but its mechanism of action remains unclear. Here, we found that, rather than acting as an instructive signal, E93 promotes adult differentiation through the repression of the pupal specifier *broad (br)*. In the absence of E93, sustained high levels of Br during the pupal stage strongly represses pupal-specific enhancers that are essential for the terminal differentiation of the wing. We also show that Br represses the pupal-enhancers during the larval and prepupal stages preventing the premature implementation of the adult genetic program, and that it also dampens the activity of larval enhancers during the latter stages of larval development. This mechanism of action seems to be a derived feature acquired in Diptera, as in the coleopteran *Tribolium castaneum*, repression of *br* by E93 is not sufficient to allow adult differentiation. In summary, our study elucidates the crucial role of the intricate interplay between E93 and Br as the governing mechanism in the process of terminal differentiation in *Drosophila*. This discovery holds significant implications for advancing our understanding of the evolution of insect metamorphosis.

## INTRODUCTION

The successful development of adult insects relies fundamentally on the precise coordination of two pivotal regulatory axes: the spatial axis, responsible for determining tissue and cell identity, and the temporal axis, governing the identity and sequential progression of diverse developmental stages throughout the life history of organisms. Recent studies have challenged conventional assumptions, revealing that the temporal transcriptional program holds more sway than the tissue-specific transcriptional program (1, 2).

In insects, the precise temporal regulation of development, encompassing the identity of each successive stage and their orderly progression, is primarily orchestrated by two hormones: 20-hydroxyecdysone (hereafter referred to as ecdysone) and Juvenile hormone (JH) (3–8). These hormones temporally regulate the expression of a specific set of genes encoding conserved stage-identity transcription factors, collectively known as the *Metamorphic Gene Network* (MGN) (9). In the holometabolous insect *Drosophila melanogaster*, the MGN encompasses three temporal factors: Chronologically inappropriate morphogenesis (Chinmo) (10, 11) and Broad (Br) (12), both members of the bric-a-brac-tramtrack-broad family, and Ecdysone inducible protein 93F (E93), a helix-turn-helix protein (13, 14). The regulatory actions of ecdysone and JH, combined with the mutual repression among *chinmo*, *br* and *E93*, ensure their precisely sequential expression throughout development (10, 11), enabling them to regulate the genetic programs governing the larval, pupal and adult stages, respectively.

While the significance of these temporal factors in delineating specific developmental windows is well established, their underlaying mechanism of action remain poorly understood. A particular important area of study involves deciphering the mechanisms of action of the adult specifier E93, given its pivotal role in controlling the terminal differentiation of adult structures during the metamorphic transition. Initially identified as an ecdysone-dependent gene essential for metamorphosis progression in *Drosophila* (15, 16), its universal role as an “adult-specifier” in all metamorphosing insects was firmly established through gene knockdown experiments in both hemimetabolous and holometabolous insects (13, 17). In *Drosophila*, E93 expression occurs during the prepupal and pupal stages, and its depletion leads to developmental arrest at the end of the pupal stage, with a significant impairment of adult differentiation (13, 18).

Recent discoveries have demonstrated the instructive role of E93 in regulating gene expression by modulating chromatin accessibility during the metamorphic pupal stage. Studies of E93-depleted wings showed that this transcription factor plays a crucial role in modulating the activity of gene enhancers. E93 activates enhancers that are active in the pupal stage and represses enhancers that were active in larval and prepupal stages, governing the timing of gene expression during wing maturation (19, 20). However, these studies were based on the reduction of E93 levels in wings during the pupal stage, and a critical aspect was not considered: the repression of *br* expression by E93 during this stage (13, 14). This repression ensures that only the appropriate temporal determinant, E93, is present during the metamorphic period. Thus, the absence of E93 in pupal wings prolongs the misexpression of Br throughout the entire pupal stage, rather than its normal disappearance (13). This critical observation prompts us to explore the extent to which the regulatory interaction between Br and E93 intricately influences the terminal differentiation of the *Drosophila* wing.

Here, we report an unexpected role of E93 in wing pupal development. Rather than acting as an instructive cue for enhancers during early and late pupal stages, the regulatory effect of E93 is mediated through repressing *br* during this period. Remarkably, our findings demonstrate that in the absence of E93, persistent Br becomes a strong repressor of the previously defined as E93-dependent pupal enhancers (19, 20). Consequently, terminal differentiation of *Drosophila* wings can proceed even without both E93 and Br once the wing has entered the pupal period. We further show that Br acts as a repressor of pupal enhancers during the larval and prepupal stages, effectively preventing premature deployment of the adult genetic program. Interestingly, we also show that Br not only prevents premature induction of pupal enhancers but also suppresses larval enhancer activities towards the end of larval development. Finally, we demonstrate that in the less derived holometabolous insect *Tribolium castaneum*, E93 is essential for adult differentiation regardless of the repression of *Br*, thus suggesting that E93 has lost its instructive role during the evolution of highly derived holometabolan insects such as *Drosophila*. Altogether, these insights shed light on the intricate regulatory network governing wing development during metamorphosis. Taken together, our analysis demonstrates that the repressive effect of E93 on *br* expression emerges as a pivotal event orchestrating the precise timing and sequential activation of enhancers across the metamorphic stage, shaping the remarkable process of terminal differentiation in *Drosophila* wings.

## RESULTS

### *Drosophila* E93 orchestrates wing terminal differentiation by repressing *br*

In *Drosophila*, the adult specifier E93 is expressed specifically throughout the prepupal and pupal stages, orchestrating the final differentiation of imaginal tissues (13, 16, 18). To investigate the specific role of E93 on wing development, we selectively depleted E93 in the wing by using the *rotund* (*rn*)-*Gal4* driver. This manipulation, mirroring the absence of E93 during larval development, had no discernible effects on wing disc development or evagination during the larval stages. However, upon entering the pupal stage, a significant impairment in terminal adult differentiation became evident (Fig. 1A and B). The wings displayed a noticeable reduction in size, along with a striking absence or poor definition of veins, and hair polarity defects (Fig. S1) (14, 20). These observations provide strong evidence for the pivotal role of E93 in driving final wing differentiation, likely achieved through the activation of genes responsible for terminal differentiation.

**Figure 1.**
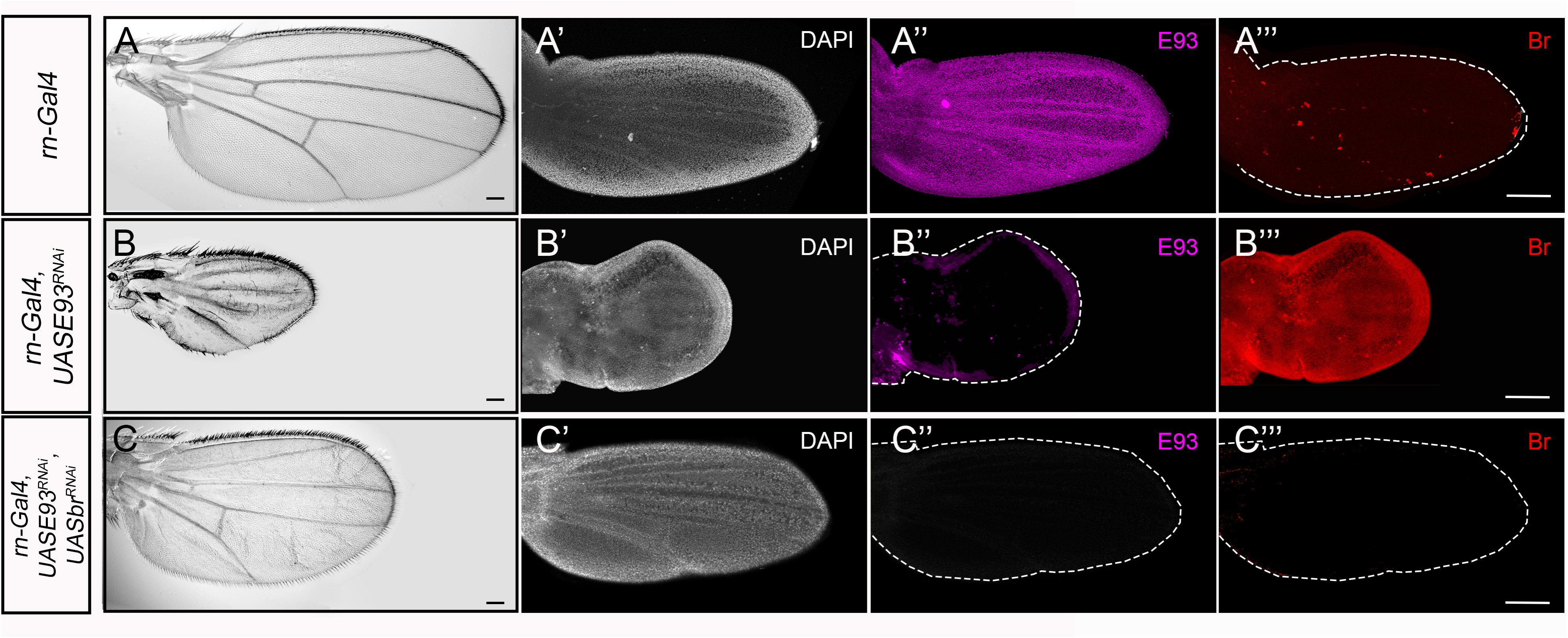
E93 controls terminal differentiation of the *Drosophila* wing by repressing *br*. (A-C) Representative images of control (A), E93-depleted (B), and simultaneously E93- and Br-depleted (C) adult wings. (A’-C’’’) Representative images of 24 h after puparium formation pupal wings of the indicated genotypes stained with DAPI to visualize nuclei (A’-C’), anti-E93 (A’’-C’’), and anti-Br (A’’’-C’’’). The white broken lines indicate pupal wing contours. Scale bars: 100 μm.

Notably, pupal wings subjected to *rn-Gal4*, *UAS-E93^RNAi^*also displayed persistently elevated levels of Br during the pupal period (Fig. 1A’-B’’’), and increased apoptosis, as indicated by elevated expression of the effector caspase Dcp-1 (Fig. S2A-C). We thus investigated whether the phenotypic consequences of E93 depletion were ascribed to the ectopic expression of Br or to the increased apoptosis of wing cells. We found that the apoptotic effect induced by E93 depletion was not responsible for the impaired differentiation of the wing. Overexpression of the baculovirus protein p35, a potent inhibitor of effector caspases (21), in E93-depleted wings did not restore normal development of the wing (Fig. S2D-G). Conversely, simultaneous depletion of *E93* and *br* in the prepupal stage onwards, using the *rn-Gal4* driver and the temperature-sensitive *Gal80^ts^* repressor system, resulted in wings undergoing normal development during the pupal period. Although smaller, these wings displayed accurate venation, well-defined hairs with proper orientation, and appropriately positioned ventral and dorsal surfaces (Fig. 1C and S2). Collectively, these findings indicate that E93 functions primarily through the suppression of *br* expression during the pupal stage, rather than acting instructively, as previously suggested (19).

### E93-dependent activation of pupal-specific enhancers is mediated by *br* repression

The above findings demonstrate that E93 primarily acts as a regulator of adult differentiation in *Drosophila* wings by suppressing *br* expression. To validate this hypothesis, we investigated the regulation of well-characterized pupal-specific enhancers crucial for wing terminal differentiation: *tnc^blade^* and *nub^vein^*. Both enhancers rely on E93 for proper activation, with their accessibility profile and activity spanning the entire wing pupal developmental stage (19, 20). Specifically, *tnc^blade^* exhibits robust activity from early to mid-pupal stages, while *nub^vein^* becomes prominent in the late-pupal stage (Fig. 2A).

**Figure 2.**
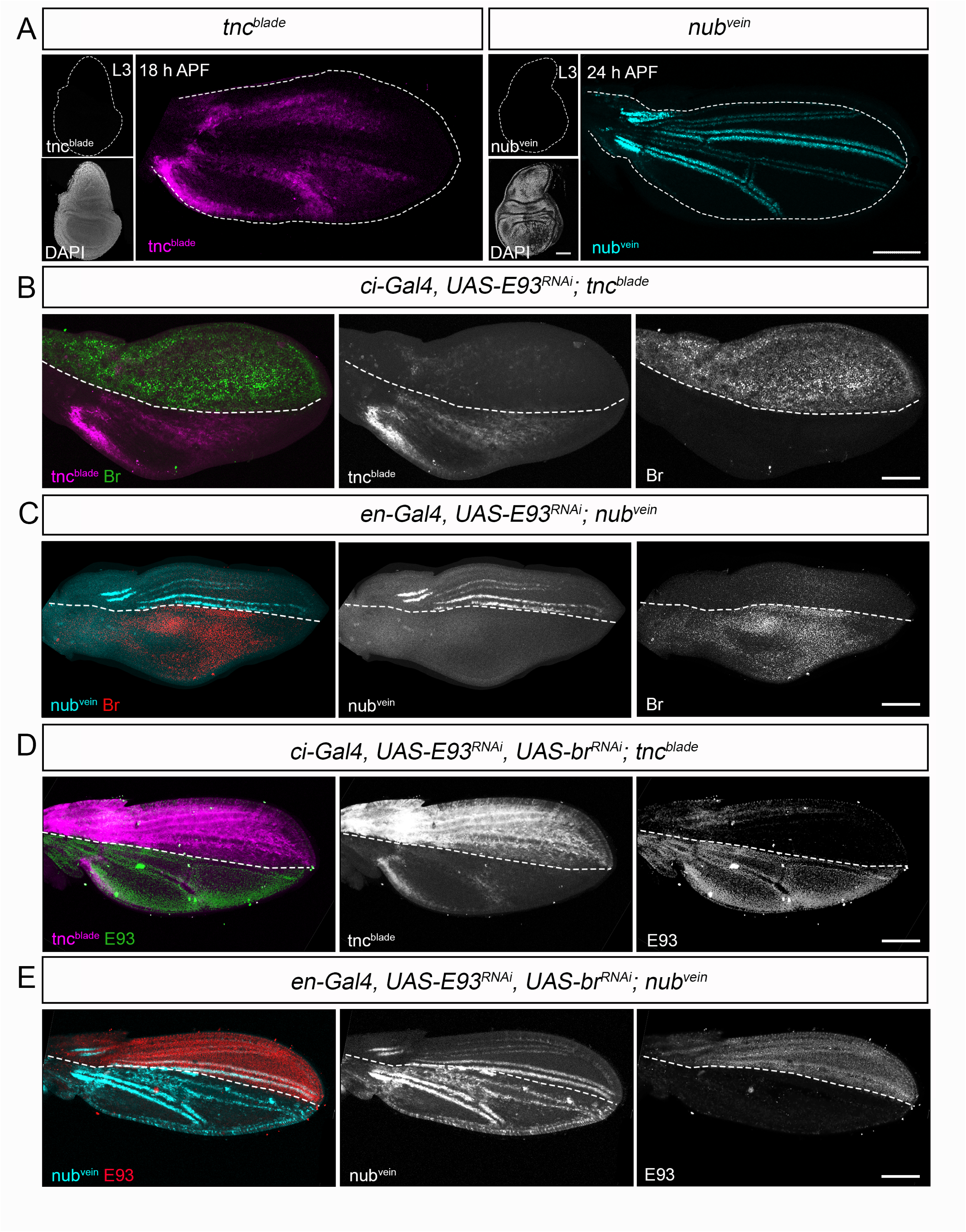
The activation of pupal-specific enhancers via E93 involves *br* repression. (A) Confocal images of *tnc^blade^* and *nub^vein^* enhancer activities. Both enhancers are inactive in wing discs and active in pupal wings. *tnc^blade^* is active (purple) in the intervein region of the wing blade while *nub^vein^* is active (blue) along longitudinal veins. Wing discs were labelled with DAPI to visualize nuclei. (B, C) Representative images of *tnc^blade^* (B) and *nub^vein^* (C) enhancer activities in pupal wings expressing *E93^RNAi^* under *control* of *ci-Gal4* or *en-Gal4*, respectively. Wings were stained with anti-Br to visualize the ectopic expression of Br in the corresponding E93-depleted regions. (D, E) Representative images of *tnc^blade^* (D) and *nub^vein^* (E) enhancer activities in pupal wings simultaneously expressing *E93^RNAi^*and *br^RNAi^* under the control of *ci-Gal4* or *en-Gal4*, respectively. Wings were stained with anti-E93 to visualize the absence of expression of E93 in the E93/Br-depleted domains. In A, wing disc and pupal wing contours are labelled by white lines. In B-E, dotted lines indicate the boundary of the corresponding driver expression domain. First panels in B-E are overlay images of the corresponding panels on the right. Scale bars: 100 μm.

As previously demonstrated (20), targeted depletion of E93 in the anterior or posterior compartments of the pupal wing, achieved through the *cubitus interruptus* (*ci*)-*Gal4* driver for *tnc^blade^* or the *engrailed* (*en*)-*Gal4* driver for *nub^vein^*, resulted in the loss of both enhancer activities in the respective compartments of early- and late-pupal wings (Fig. 2B and C). As before, the absence of E93 also led to sustained high levels of Br in both cases (Fig. 2B and C). In line with our previous observations, simultaneous depletion of both *E93* and *br* after wing eversion remarkably reinstated the activities of *tnc^blade^* and *nub^vein^*in the corresponding compartments (Fig. 2D and E and S3). These data firmly establish that (i) E93 binding to *tnc^blade^*and *nub^vein^* is not a prerequisite for their induction in pupal wings and (ii) affirm that the regulatory effect of E93 during wing pupal development primarily operates through the repression of *br*.

### Br represses pupal-specific enhancers in larval-prepupal wings

Thus far, we have presented compelling evidence suggesting that Br has the capacity to suppress pupal-specific enhancers during the pupal stages. Nevertheless, a previous study demonstrated that overexpression of E93 in wandering L3 (L3W) or early prepupal wings leads to the premature expression of *tnc^blade^* and *nub^vein^* enhancers, supporting the instructive role of E93 (19). However, in line with our findings, we found that the early activation of *tnc^blade^* (*ci-Gal4, UAS-E93*) or *nub^vein^* (*en-Gal4, UAS-E93*) enhancers was concurrent with a significant repression of *br* in both cases (Fig. 3A and C). This strongly supports that the absence of Br triggers the precocious activation of pupal enhancers in larval and prepupal stages.

**Figure 3.**
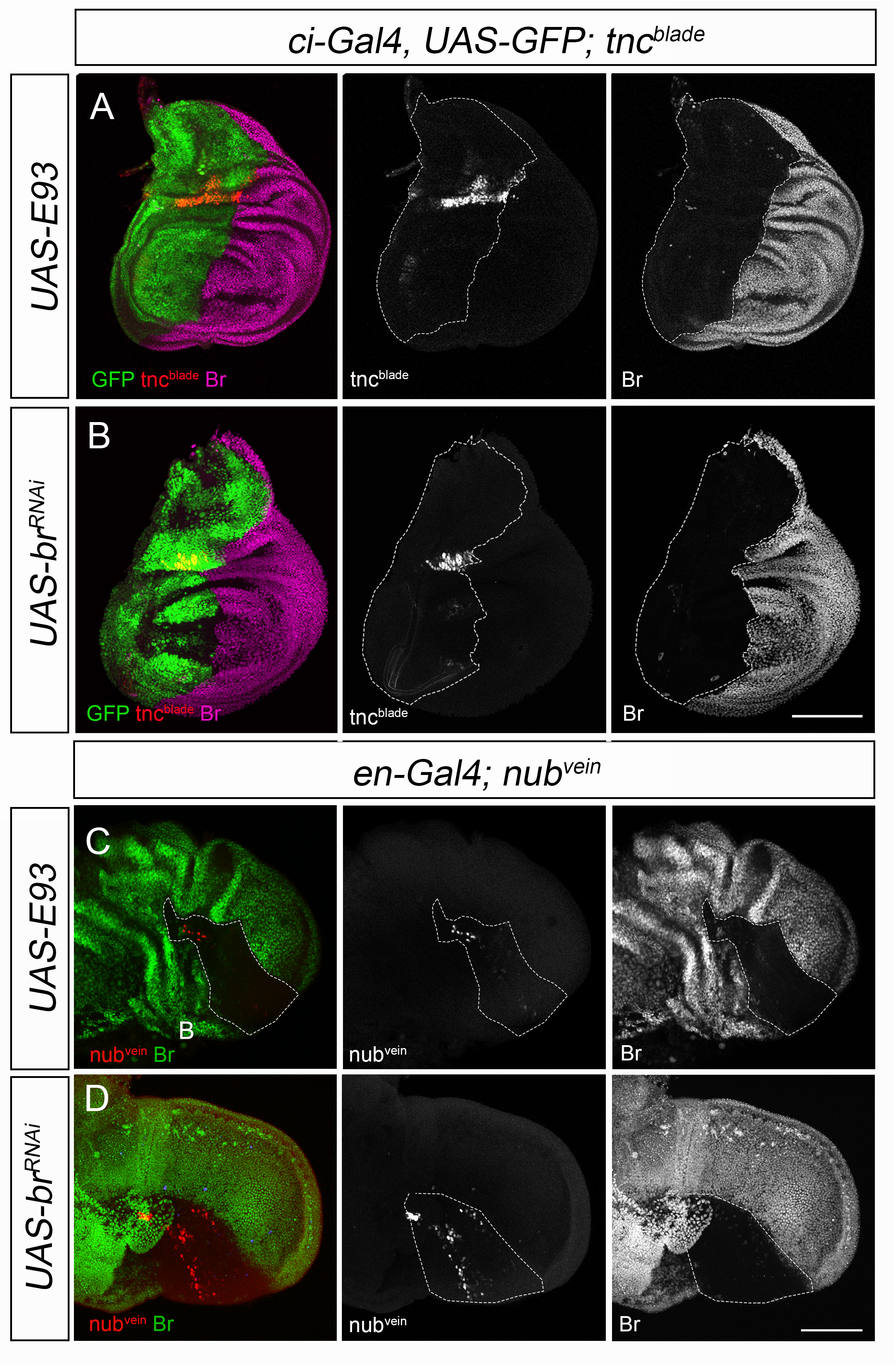
Br suppresses pupal-enhancers in larval wing discs and prepupal wings. (A, B) Confocal images of *tnc^blade^* enhancer activity and immunostaining of Br in wandering L3 wing discs expressing ectopic *E93* (A) or *br^RNAi^* (B) under the control of *ci-Gal4*. (C, D) Confocal images of *nub^vein^* enhancer activity and immunostaining of Br in 9h prepupal wings expressing ectopic *E93* (C) or *br^RNAi^* (D) under the control of *en-Gal4*. Dotted lines indicate the boundary of the corresponding driver expression domain. First panels are overlay images of the corresponding panels on the right. Scale bars: 100 μm.

To confirm that the absence of Br, rather than the premature presence of E93, was responsible for the precocious activation of both enhancers, we selectively depleted *br* in either the anterior or posterior compartment of larval wing discs (*ci-Gal4, UAS-br^RNAi^*) and prepupal wings (*en-Gal4, UAS-br^RNAi^*), respectively. Notably, while the absence of Br did not lead to a precocious up-regulation of E93 (Fig. S4), we observed a robust premature activation of both *tnc^blade^* and *nub^vein^* in their respective regions (Fig. 3B and D and S4). Altogether, these results demonstrate that these pupal-specific enhancers can be activated during the larval-prepupal stages, and that their activity remains suppressed due to the repressive influence of Br during this critical transitional period.

### The Chinmo-to-Br switch prevents the precocious ecdysone-dependent *tnc^blade^* activity in larval wing discs

To determine precisely the developmental stage at which late pupal enhancers can become active, we investigated the activity of *tnc^blade^*across various larval developmental stages using *ci-Gal4, UAS-br^RNAi^*in wing discs. Under these conditions, we detected *tnc^blade^* activity exclusively from mid-L3 wing discs onwards (Fig. 4A-C). Notably, this stage marks the transition of wing disc cells from a state refractory to differentiation to one permissive for differentiation. This transition is governed by the reciprocal repression between the larval specifier Chinmo and Br (10, 11, 22).

**Figure 4.**
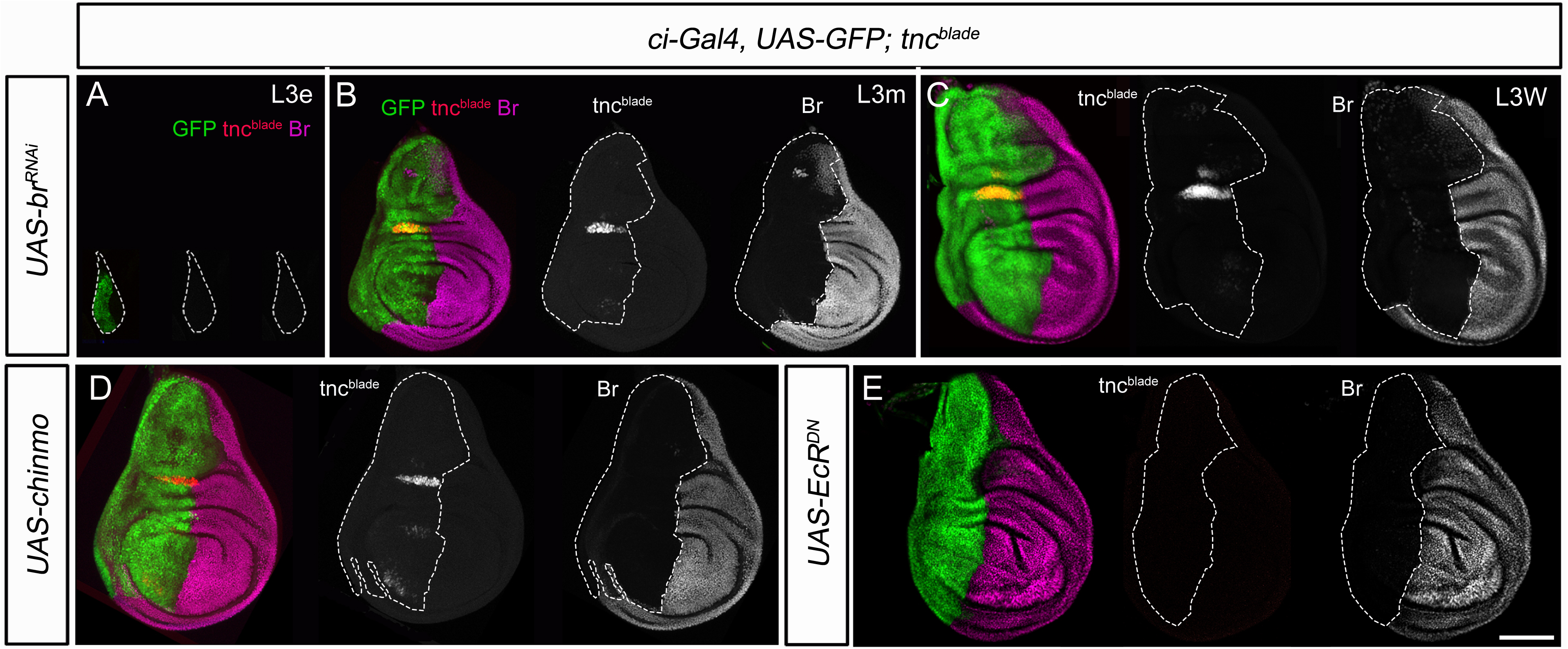
The Chinmo-to-Br switch at mid-L3 prevents the premature activation of *tnc^blade^* by ecdysone. (A-C) Confocal images of *tnc^blade^* enhancer activity and immunostaining of Br in early L3 (eL3) (A), mid-L3 (mL3) (B) and wandering L3 (WL3) (C) wing discs expressing *br^RNAi^* under the control of *ci-Gal4*. (D) Confocal images of *tnc^blade^* enhancer activity and immunostaining of Br in WL3 wing discs expressing chinmo under the control of *ci-Gal4*. (E) Confocal images of *tnc^blade^* enhancer activity and immunostaining of Br in L3W wing discs expressing the dominant negative form of *EcR* (*EcR^DN^*) under the control of *ci-Gal4*. In all panels, dotted lines indicate the boundary of the corresponding driver expression domain. First wing discs in all panels are overlay images of the corresponding panels on the right. Scale bars: 100 μm.

Given that Br represses *tnc^blade^* pupal enhancer activity from mid L3 onwards, we considered whether the larval specifier Chinmo played a crucial role in maintaining the enhancer in a repressed state during early larval development. To explore this hypothesis, we examined the impact of sustained high levels of Chinmo in L3W wing discs on *tnc^blade^* enhancer activity. Notably, overexpression of Chinmo in the anterior compartment of the disc starting in early L3 suppressed *br* expression but failed to repress pupal enhancers, as evidenced by a clear premature activation of *tnc^blade^* activity in L3W wing discs (Fig. 4D). These findings demonstrate that the induction of *tnc^blade^* activity occurs at the Chinmo-to-Br switch around mid-L3, but is actively prevented by the repressive function of Br.

To explore the signal involved in the induction of *tnc^blade^*activity which starts in mid-L3, we studied whether increased ecdysone signaling following the Chinmo-to-Br transition is sufficient for enhancer activation. For this, we overexpressed a dominant negative form of the ecdysone receptor, EcR^DN^, in the anterior compartment of the disc using *ci-Gal4*. EcR^DN^ functions by preventing the binding of its ligand 20E, thereby inhibiting the activation of the ecdysone signaling pathway (23). Surprisingly, despite the complete absence of Br expression after EcR^DN^ overexpression, we did not observe premature induction of *tnc^blade^* activity in this compartment (Fig. 4E). This discovery strongly suggests that ecdysone signaling subsequent to the Chinmo-to-Br transition is necessary for triggering the activation of this enhancer.

### Larval enhancers are also repressed by Br

So far, we have demonstrated that Br functions as a temporal regulator ensuring that pupal enhancers do not activate prematurely. This suggests the possibility that Br could also terminate the larval genetic program at the end of juvenile development by repressing the activity of larval enhancers. To study this possibility, we aim to investigate the role of Br in regulating the activity of early larval enhancers using *wingless^1^*-reporter (*wg^1^-lacZ*) as the experimental model. *wg1* is a highly conserved enhancer of the *wg* gene responsible for specifying wing fate in the developing wing primordium (24). The activity of *wg^1^-lacZ* closely mirrors the dynamic expression pattern of Wg in the ventral anterior compartment of the wing disc during early larval stages, with a significant decrease in enhancer activity by mid-late L3, coinciding with the occurrence of Br (Fig. 5A) (25).

**Figure 5.**
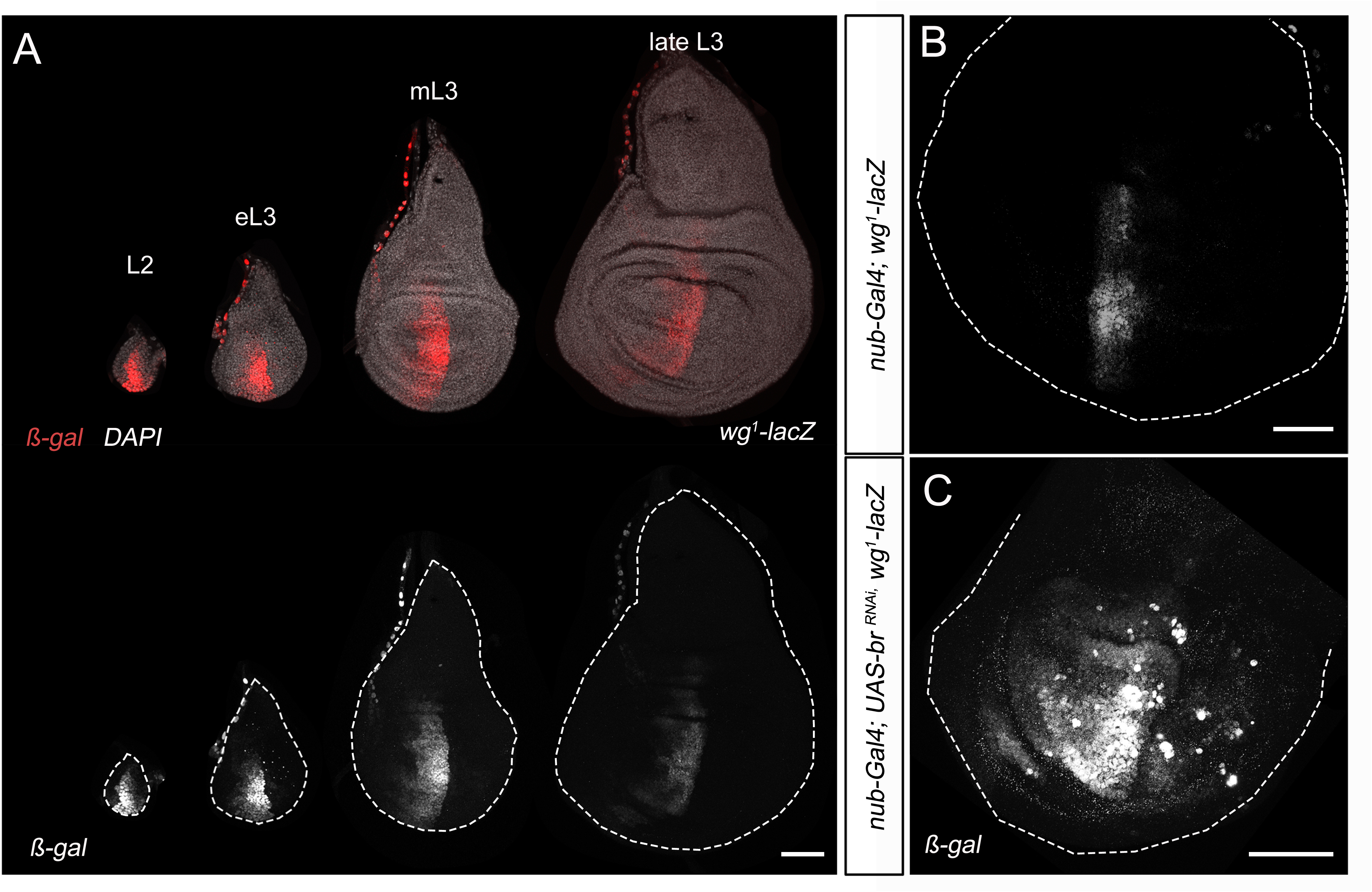
Br represses the larval *wg^1^* enhancer. (A) Confocal images of second (L2), early (eL3), mid (mL3) and late third (late L3) instar wing discs of larvae bearing the *wg^1^-lacZ* reporter construct and stained for ß-galactosidase. Wing discs are stained with DAPI to visualize nuclei. (B) Reporter activity for the *wg^1^-lacZ* reporter construct in WL3 wing discs expressing *br^RNAi^* under control of *nub-Gal4.* Wing disc contours are labelled by white lines. Scale bars: 50μm.

To investigate the potential role of Br in regulating the decline of *wg^1^-lacZ* activity during late L3, we depleted *br* expression in the wing disc pouch using the *nubbin (nub)-Gal4* driver. Remarkably, we observed that the activity of *wg^1^-lacZ* was not correctly suppressed but instead, exhibited a significant expansion in the anterior compartment (Fig. 5B and C). Altogether, these findings demonstrate that Br plays a critical role as a developmental timer at the larval-pupal transition by repressing larval enhancers and, at the same time, avoiding the premature activation of late-pupal enhancers.

### E93-dependent terminal differentiation does not depend on *br* repression in less derived holometabolans

Finally, we investigated the conservation of the E93 regulatory mechanism in terminal differentiation across holometabolous insects. To address this, we turned to the basal holometabolous insect, *Tribolium castaneum*, where the functions of E93 as an adult specifier and its ability to suppress *br* expression during the pupal stage have been firmly established (14, 17). First, we specifically targeted E93 depletion during the pupal period using RNA interference (RNAi) by injecting double-stranded RNA (*dsRNA*) against *E93* in early pupae. Consistent with previous findings (14), *E93-*depleted pupae did not exhibit any sign of adult differentiation by the end of the pupal period (Fig. 6A). For example, metamorphic changes in the legs, such as segmentation, pigmentation, and significant alterations in shape, which typically occur during the pupal stage, were entirely prevented (Fig. 6B). Confirming that the adult genetic program has not been installed in the E93-depleted pupae, *br* expression was not properly repressed and the activation of the adult-specific cuticle gene *Cpr27*, which specifically occurs during the pupal–adult molt (26), was completely eliminated (Fig. 6C).

**Figure 6.**
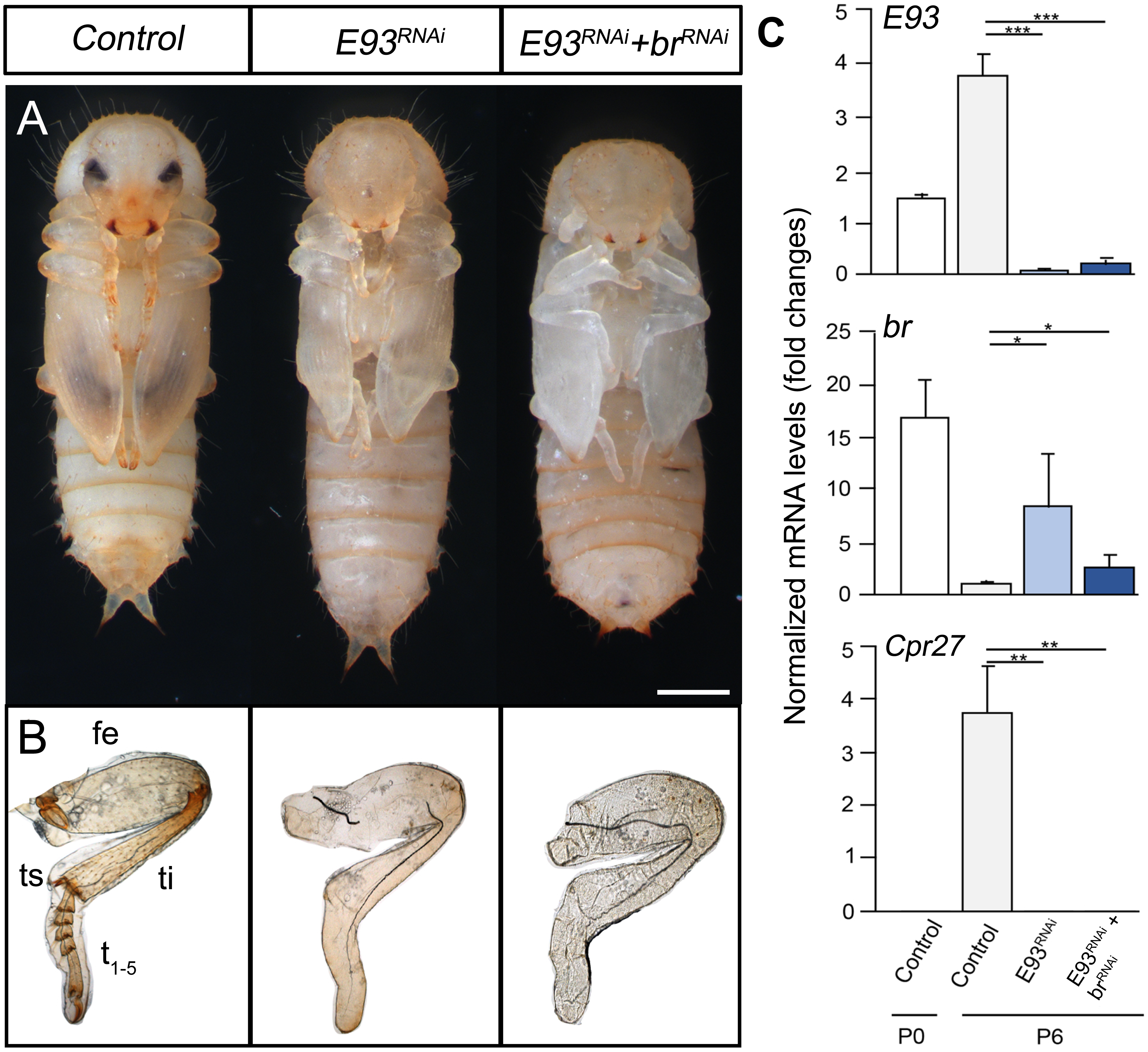
Adult differentiation in the basal holometabolous *Tribolium castaneum* is controlled by E93 but is not channeled through *br* repression. (A) Ventral views of 6-day-old *Control*, *E93^RNAi^*, and *E93^RNAi^+br^RNAi^* pupae. Newly molted pupae were injected with *dsMock* (*Control*), *dsE93* (*E93^RNAi^*), or *dsE93* + *dsbr* (*E93^RNAi^+br^RNAi^*) and left for six days. (B) Comparison of the external morphology of the mesothoracic leg of 6-day-old *Control*, *E93^RNAi^*, and *E93^RNAi^+br^RNAi^*pupae. Abbreviations: fe, femur; ti, tibia; ts, tibial spur; t1-5, tarsomeres 1-5. (C) Transcript levels of *E93*, *br*, and the adult-specific gene *Cpr27* were measured by qRT-PCR in 0-day-old (P0) *Control* pupa, and 6-day-old (P6) *Control*, *E93^RNAi^*, and *E93^RNAi^+br^RNAi^*pupae. Transcript abundance values are normalized against the *RpL32* transcript. Error bars indicate the SEM (n = 3-5). Asterisks indicate differences statistically significant at *\*p*≤0.1; *\*\*p*≤0.01 and *\*\*\*p*≤0.001 (*t*-test). Scale bar: 0,5 mm.

Next, to investigate the impact of elevated *br* levels during the pupal stage, we simultaneously depleted *E93* and *br* by injecting the corresponding *dsRNAs* into newly molted pupae. These pupae displayed no signs of adult differentiation (Fig. 6A), with their appendages maintaining a typical pupal-like morphology (Fig. 6B). Consistent with this phenotype, the expression of *Cpr27* was completely suppressed in the double knock-down animals, confirming that, in contrast to *Drosophila*, the adult genetic program has not been initiated (Fig. 6C). These findings indicate that in less derived holometabolous insects, terminal differentiation controlled by E93 does not occur through the repression of *br*, thus suggesting the likely loss of the instructive role of E93 in more derived holometabolous insects, such as *Drosophila*.

## DISCUSSION

Adult differentiation in *Drosophila*, as observed in other holometabolous insects studied thus far, is orchestrated by the master adult-specifier factor E93 during the pupal stage (13, 14, 27–35). Recent genomic studies have revealed that E93 plays a crucial temporal role in terminal differentiation by orchestrating chromatin dynamics. This orchestration involves opening gene enhancers critical for terminal differentiation while simultaneously silencing those active in preceding stages (19, 20). However, our results demonstrate that E93 does not function as an instructive factor for terminal differentiation. Instead, our findings indicate that E93 operates through the repression of *br* expression during the pupal stage. This repression holds paramount significance, as Br acts as a potent suppressor of pupal enhancer activities when expressed during the pupal period (Fig 2B and C). Our results demonstrate that eliminating the Br-mediated repression on pupal-specific enhancers is sufficient to activate them, regardless of E93 presence (Fig 2D and E and S3). This finding is further supported by the emergence of fully developed adult wings that differentiate in the absence of both E93 and Broad throughout the pupal stage (Fig 1 and S1B).

Does this regulatory mechanism extend beyond these enhancers, having broader implications for adult wing differentiation? While our genetic analysis focused on two specific E93-dependent enhancers, *tnc^blade^*and *nub^vein^*, it is important to emphasize that these enhancers exemplify a broader class of enhancers active during the prepupal and pupal stages that exhibit significant enrichment of Br binding sites (19), which suggests that Br functions as a pervasive temporal repressor for pupal-specific enhancers. However, whether Br modulates accessibility of those enhancers or adopts a classical transcriptional repressor role upon binding to accessible enhancers warrants further investigation. Despite this uncertainty, the downregulation of *br* by E93 during the pupal stage emerges as the pivotal mechanism governing terminal differentiation, as it enables the proper temporal activation of pupal-specific enhancers.

Importantly, the repressive effect of Br on these pupal enhancers holds significant importance for fine-tuning the temporal orchestration of molecular events underlying the larval-to-pupa transition. We have demonstrated that this regulatory mechanism maintains pupal-enhancers in an inactive state during the period of heightened *br* expression (Fig 3B and D and 4A-C) and operates independently of E93 (Fig 3 and S3). This observation strongly supports the previously stated concept that E93 does not serve as an instructive signal for activating these enhancers during the pupal period.

On the other hand, our results regarding the distinct timings of premature induction for *tnc^blade^* (beginning in mid-late L3) and *nub^vein^* (beginning in the prepupal stages) in wings lacking Br underscore the gradual emergence, from mid-L3 onwards, of the combination of developmental inputs required for activating these enhancers. Consequently, the presence of Br, from mid-L3 until the end of the prepupal stage, prevents the premature initiation of the terminal differentiation program, ensuring that the undifferentiated wing disc has sufficient time to fully grow and undergo eversion before entering the final phases of terminal differentiation. In this context, our findings unveil that the acquisition of enhancer competence for induction from mid-L3 stems from the increasing levels of ecdysone, which occurs when the larva attains the “critical weight” (CW) checkpoint - a developmental growth-related milestone marking the irreversible commitment to metamorphosis (36). Notably, the rise in ecdysone levels also triggers the transition from *chinmo* to *br* expression, delineating the cellular shift from a state of differentiation resistance to a state of differentiation permissiveness (10, 11, 22). Thus, the surge in ecdysone signaling post-CW exerts a dual effect: it instructs enhancer inducibility while simultaneously triggering *br* expression, thereby preventing the untimely activation of such enhancers.

In summary, we conceive a model in which a robust surge in ecdysone levels following the CW checkpoint acts as a catalyst for initiating the metamorphic transition into adulthood. This surge triggers the activation of enhancers linked to the expression of genes essential for final adult differentiation. However, this transition is momentarily delayed due to the presence of Br, an ecdysone-dependent gene itself, thereby maintaining the dormancy of these enhancers during metamorphosis initiation and setting the stage for progression to the intermediate pupal stage. Following the eversion of the imaginal discs, the heightened expression of *E93* at the onset of pupal period leads to *br* repression, thereby allowing reactivation of pupal-specific enhancers and ultimately facilitating the ultimate differentiation of the organism toward its terminal state (Fig. 7).

**Figure 7.**
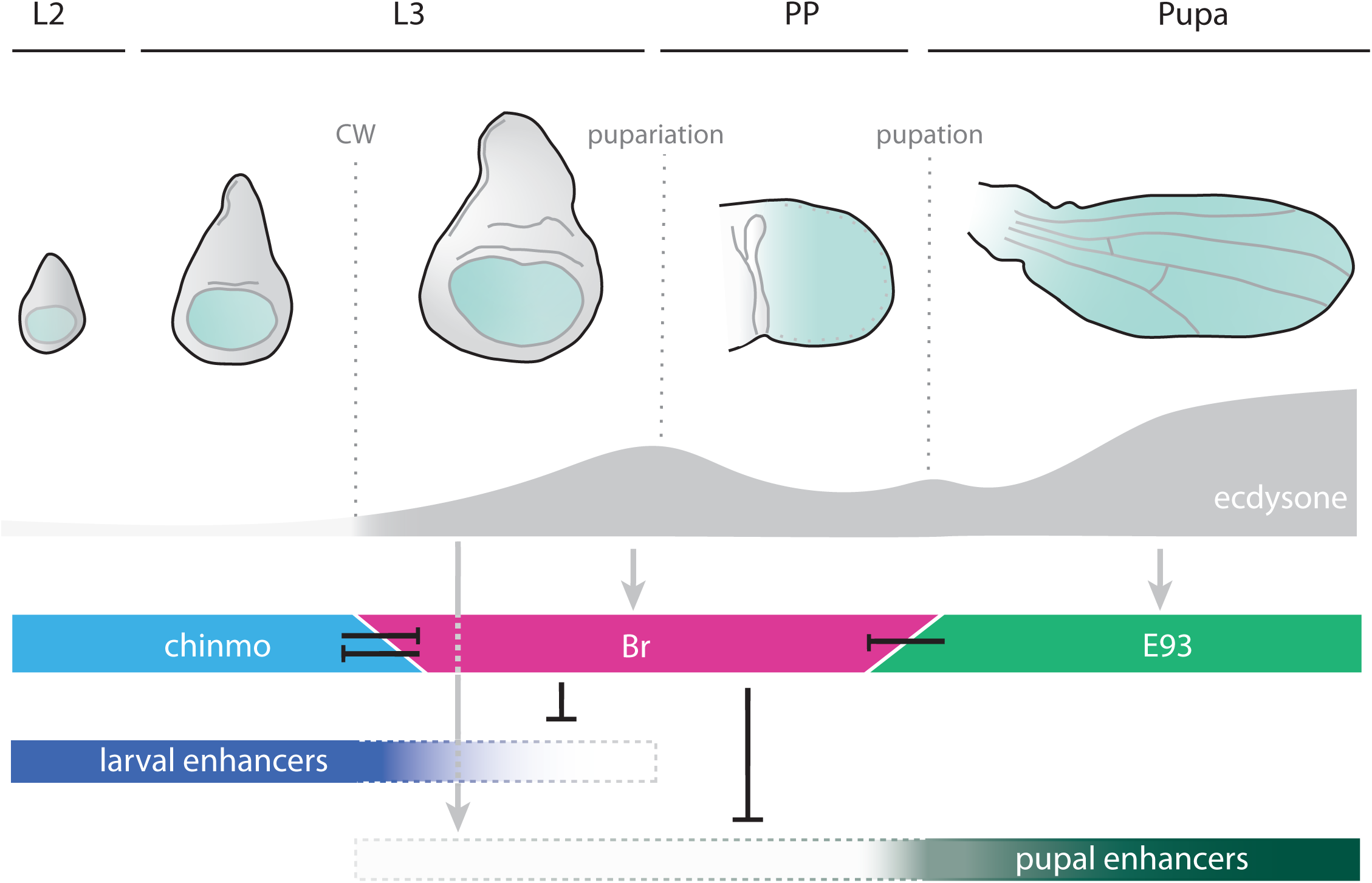
Model depicting the regulatory activities of E93 and Br in controlling adult differentiation of *Drosophila* wings during development. Ecdysone levels (grey), and stage-specific expression of chinmo (blue), Br (pink) and E93 (green) during *Drosophila* development. The critical weight (CW) checkpoint at early-mid L3 stage marks the *chinmo*-to-*broad* expression switch. Black lines represent repressive effects between chinmo, Br and E93, as well as the repressive effects of Br on the activity of larval (dark blue) and pupal (dark green) enhancers. Grey lines represent inductive effects mediated by ecdysone-dependent signaling.

Significantly, our findings not only establish the pivotal role of Br in suppressing the activity of pupal-enhancers linked to terminal differentiation during the larval-to-pupal transition, but also underscore its necessity in repressing larval-enhancers (as demonstrated in the case of the *wg^1^* enhancer). Although responsible for directing the characteristic expression of Wg in the wing disc to dictate wing fate specification during early larval phases, the *wg^1^* enhancer experiences a substantial reduction in activity as the organism reaches the mid-late L3 stage (24, 25). Our study reveals that this decline in *wg^1^* enhancer activity during the later stages of larval development also depends on Br (Fig 5B and C). The repression exerted by Br on early-larval enhancers and its role in terminating the juvenile developmental program, consequently inducing a permissive state for differentiation in wing cells, holds significance not only for developmental progression but also bears crucial implications for regeneration and tumor progression. Our findings, along with others (11, 22), strongly indicate that the regenerative capacity of damaged wing larval cells is contingent upon an environment lacking Br, thereby sustaining wing cells in a differentiation-refractory state. Indeed, it has been demonstrated that the *wg^1^* enhancer is activated in wing disc cells upon tissue injury, contributing to wing regeneration (25, 37), and further, this enhancer is repressed by Br in a context of tissue damage (38).

These observations highlight the dual role played by Br in orchestrating the temporal regulation of stage-specific genetic programs. On the one hand, it acts as a suppressor of larval-enhancers, simultaneously repressing the larval-specifier factor Chinmo, thereby marking the end of the larval developmental phase. On the other hand, it prevents premature activation of late pupal-enhancers, avoiding a premature and detrimental direct metamorphic transition to the adult form (Fig. 7). In this context, Br can be likened to a temporal hinge, facilitating a seamless transition from larval to adult genetic programming. The dual repressive role of Br on both larval and adult enhancer activity aligns with its well-established role in more basal holometabolous species. For example, depletion of Br in *Tribolium* disrupts normal larva-pupal transition, leading to a terminal molt characterized by the simultaneous presence of larval and adult features in different body parts (14, 39–41). Remarkably, while the ability to repress larval genes appears to be a feature acquired in holometabolous insects, the capacity of Br to suppress the expression of adult genes may predate the evolution of holometaboly. This notion is supported by findings in the ancestral hemimetabolan insect, the odonate *Ischnura senegalensis*, where Br has been identified as capable of inhibiting a set of adult-specific genes associated with pigmentation during premetamorphic stages (29).

Finally, our results uncover significant differences between the mechanism of action of E93 as the regulator of adult terminal differentiation between highly derived (*Drosophila*) and more basal (*Tribolium*) holometabolous insects. In the case of *Tribolium*, our findings show that E93 governs adult differentiation during the pupal stage although not through the repression of *br*. Interestingly, in this beetle, E93 not only functions as the adult specifier during the pupal phase but also acts as the trigger for metamorphosis at the larval-pupal transition (17). In contrast, in *Drosophila*, this metamorphosis-triggering role has been assumed by increasing levels of Br during the mid-late stages of the final larval instar (12). We propose that the shift from E93 to Br as the catalyst for triggering the metamorphic transition in highly derived holometabolous insects may be associated with a more permissive role of E93 in adult differentiation through *br* repression.

In this context, we propose a hypothesis suggesting that the functional loss of E93 as the metamorphosis-triggering factor in *Drosophila* may be correlated with the growth of imaginal discs – groups of cells that give rise to adult structures-in the presence of JH. In the majority of insects, JH inhibits the development of adult structures during the larval or juvenile stages. Thus, *Tribolium* initiates wing development at the onset of the last larval instar upon JH decay, which is necessary for the up-regulation of *E93* expression. In *Drosophila*, however, imaginal cells start growing during the larval period in the presence of JH and the absence of E93. This continuous growth of imaginal tissues during larval development in *Drosophila* suggests an evolutionary scenario wherein E93 may have relinquished its role as a metamorphic trigger in favor of Br. Interestingly, some lepidopteran species present a mix of cell primordia and imaginal discs. Further investigations into the mechanism of action, at the genetic and genomic level, of E93 in these species would be highly valuable for understanding the evolution of the mechanism of action of E93 according with the type of developmental strategy based on the presence of imaginal discs.

## METHODS

### *Drosophila* culture and genetics

All fly stocks were cultured at 25°C on standard flour/agar *Drosophila* media. The Gal4/UAS system was employed to induce transgene expression at 29°C, and conditional activation of either RNAi or gene expression was achieved using the Gal4/Gal80^ts^ system. Experiments involving larval and prepupal stages entailed maintaining crosses at 18°C until the third larval instar, followed by a shift to 29°C 48 hours before tissue dissection. Wandering larvae were gathered for wing disc dissection, and mid-prepupa specimens were collected for prepupal wing dissection. For pupal and adult wings collection, the crossed were kept at 18°C until 0h after puparium formation (APF) and then switched to 29°C for conditional induction, with tissues dissected at the indicated time points. Experimental and control crosses were conducted in parallel.

The following lines were used in this study: *UAS-br^RNAi^* (#33641; #27272); *UAS-p35* (#5072; #5073); *UAS-chinmo* (#50740); *UAS-EcrB1^DN^* (#6872); *Tub-Gal80^ts^* (#7018, #7019); and *UAS-mCD8::GFP* (#32186) from the Bloomington Drosophila Stock Center (BDSC) and *UAS-E93^RNAi^* (KK108140) from the Vienna Drosophila RNAi Center. The *UAS-E93* line (16) was generously provided by Ian Duncan, reporter lines *tnc^blade^-tdTomato* and nub^vein^-nlsGFP (20) were provided by Daniel J. McKay, and *wg^1^-lacZ* (25) by Marco Milán. To drive the expression of different constructs in the wing disc, the *rn-Gal4* (42), *en-Gal4* (BDSC #30564), *nub-Gal4* (BDSC #86108) and *ci-Gal4* (43) lines were employed. Crosses to tthese lines to the yw line were used as controls.

### Tribolium castaneum

The enhancer-trap line *pu11* of *Tribolium* (obtained from Y. Tomoyasu, Miami University, Oxford, OH) was reared on an organic wheat flour diet supplemented with 5% nutritional yeast and maintained at a constant temperature of 29 °C in complete darkness.

### Microscopy, histological analysis and immunocytochemistry

For fluorescent imaging, dissected wing discs or pupal wings at specified time points were fixed in 4% formaldehyde. Following fixation, tissues underwent rinsing in 0.1% Triton X-100 (PBST) and were subjected to overnight incubation with primary antibodies, appropriately diluted in PBST. Subsequently, the samples were thoroughly washed with PBST and exposed to secondary antibodies (Alexa-conjugated dyes 555, 647, 633, Life Technologies, 1:500) for 2 hours at room temperature. After this incubation, tissues were extensively rinsed in PBS and mounted in Vectashield medium with DAPI (Vector Laboratories, H2000). The primary antibodies used in this study, along with their respective dilutions, include: mouse monoclonal anti-Br core (1:250, #25E9.D7) and monoclonal mouse anti-ßGal (1:200, #2314509) from the Developmental Studies Hybridoma Bank, rabbit anti-cleaved Dcp-1 (1:100, #9578) from Cell Signaling, and polyclonal rabbit anti-E93 (11). Images were captured using a Leica TCS SP5 confocal microscope and processed with either Fiji or Photoshop CC (Adobe). For bright field microscopy, wings of indicated phenotypes were dehydrated in ethanol and directly mounted in Hoyer’s solution for microscopic examination. Samples were examined with AxioImager.Z1 (ApoTome 213 System, Zeiss) microscope and images were subsequently processed using Photoshop CC (Adobe).

### Quantitative Real-Time Reverse Transcriptase Polymerase Chain Reaction (qRT-PCR)

Total RNA from staged *Tribolium* pupae was extracted using the GenEluteTM Mammalian Total RNA kit (Sigma). cDNA synthesis was performed following the previously described methods (44, 45). For quantitative real-time PCR (qPCR), Power SYBR Green PCR Mastermix (Applied Biosystems) was used to determine relative transcript levels. To standardize the qPCR inputs, a master mix containing Power SYBR Green PCR Mastermix and forward and reverse primers was prepared, with each primer at a final concentration of 100 µM. The qPCR experiments were conducted with an equal quantity of tissue equivalent input for all treatments, and each sample was run in duplicate using 2 µl of cDNA per reaction. As a reference, the same cDNAs were subjected to qRT-PCR using a primer pair specific for *Tribolium* Ribosomal *Rpl32*. All samples were analyzed on the iCycler iQReal Time PCR Detection System (Bio-Rad). The primer sequences used for qPCR for *Tribolium*:

*E93-F*: 5′-CTCTCGAAAACTCGGTTCTAAACA-3′

*E93-R*: 5′-TTTGGGTTTGGGTGCTGCCGAATT-3′

*Br-C-F*: 5′-TCGTTTCTCAAGACGGCTGAAGTG*-*3′

*Br-C-R*: 5′*-*CTCCACTAACTTCTCGGTGAAGCT-3′

*CPR27-F*: 5’-AGGTTACGGCCATCATCACTTGGA-3’

*CPR27-R*: 5’-ATTGGTGGTGGAAGTCATGGGTGT-3’

*Rpl32-F*: 5′-CAGGCACCAGTCTGACCGTTATG*-*3′

*Rpl32-R*: 5′*-*CATGTGCTTCGTTTTGGCATTGGA-3′

### Pupal RNAi injection

RNAi *in vivo* in *Tribolium* was performed as previously described (13, 14). The control dsRNA used a non-coding sequence from the pSTBlue-1 vector (dsMock). For pupal injections, a concentration of 1 µg/µl dsRNA was injected to newly molted pupae. In cases of experiments with two dsRNAs, equal volumes of each dsRNA solution were mixed and applied in a single injection. The primer sequences used to generate templates via PCR for transcription of the dsRNAs for *Tribolium* E93 and Br were:

*dsE93-F*: 5’-AAATAACGGTGATACAGTGTCAAG-3’

*dsE93-R*: 5’-TTGTAGTCCATCTCGGAGATGGAA-3’

*dsBr-F*: 5’-CAATTACCAAAGCAGCATCACATC-3’

*dsBr-R*: 5’-GGCTTTGTACTTGCGCCAACTGTT-3’

*Tribolium* dissections were carried out in Ringer’s saline and the hindlegs were mounted directly in Glycerol 70% and examined with an AxioImager.Z1 microscope.

## ACKNOWLEDGEMENTS

This project is supported by grants PGC2018-098427-B-I00 and PID2021-125661NB-100 to D.M., and X.F-M. and by grant 2017-SGR 1030 and 2021-SGR 0417 to D.M. and X.F-M funded by the Catalan Government. The research has also benefited from ERDF “A way of making Europe to D.M. and X.F-M”. We thank Carolina Reisenman for critical reading of the manuscript

