## Supplemental information for "E93 controls adult differentiation by repressing *broad* in *Drosophila*"

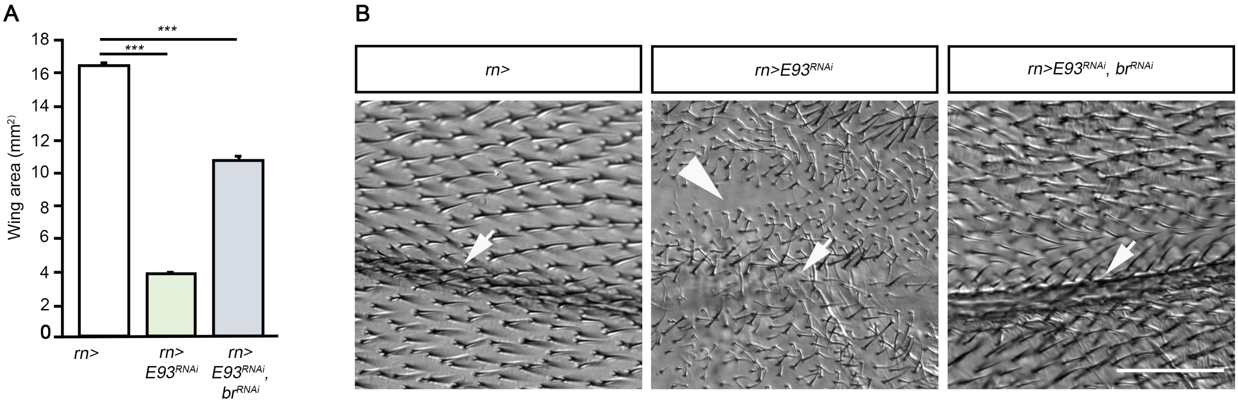


**Figure S1. Adult differentiation of *Drosophila* wing by E93 is mediated by the repression of *br*.** (A) Adult wing area measurements comparing Control (*rn*>), E93-depleted (*rn*>*E93*$^{RNAi}$) and simultaneously E93/br-depleted wings (*rn*>*E93*$^{RNAi}$, *br*$^{RNAi}$). (B) Representative image sections from adult wings of the indicated genotypes, illustrating wing hairs and venation phenotypes. Arrows denote the second wing vein and arrowhead highlight the absence of wing hairs in some areas of the E93-depleted wings. Error bars indicate the SEM (n = 7-12). Asterisks indicate differences statistically significant at ***$p \leq 0.001$ (*t*-test). Scale bar: 50 μm.

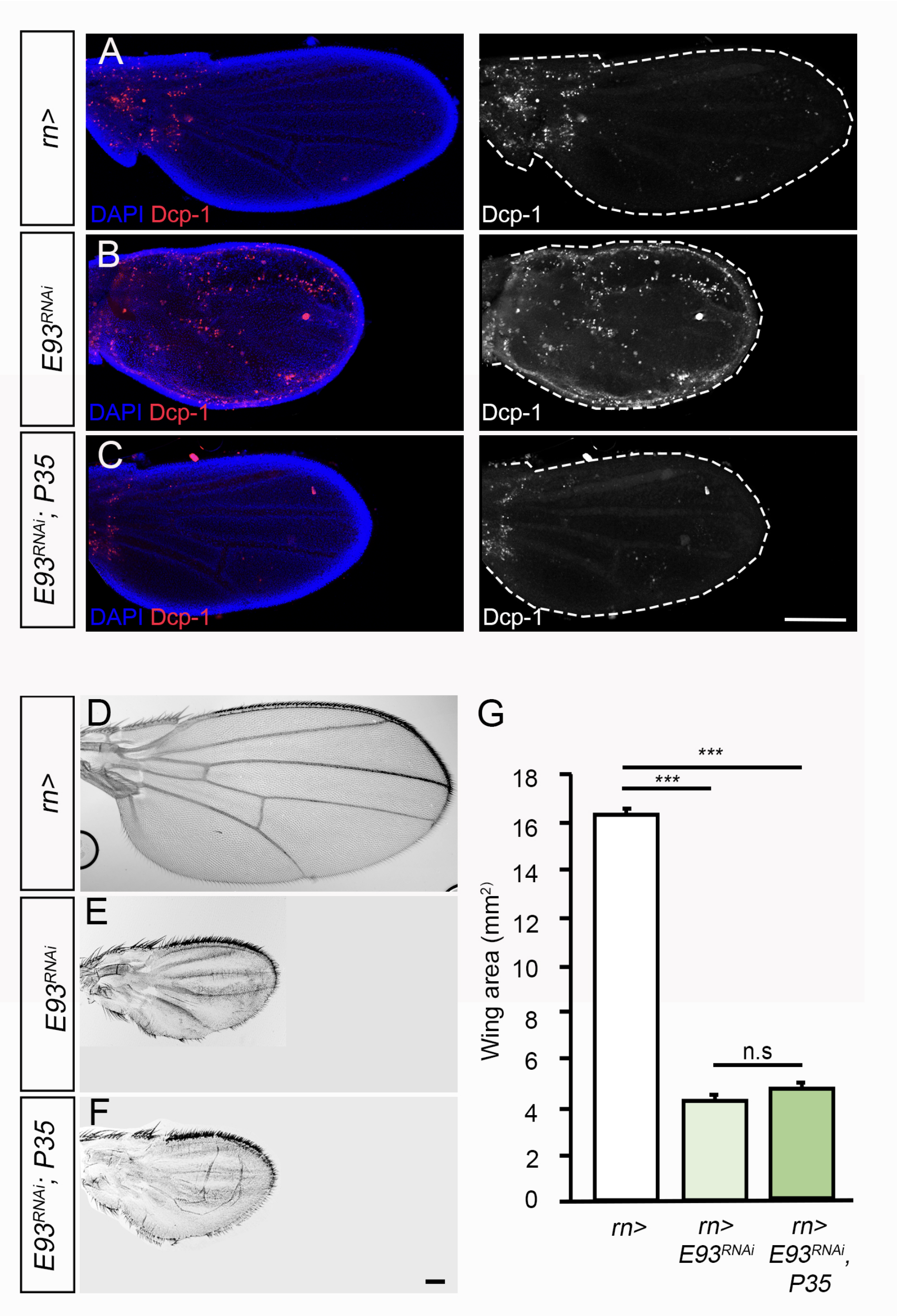


**Figure S2. E93-dependent increase in wing cell apoptosis is not responsible for the impairment of terminal wing differentiation.** (A-C) Representative images of *control* (*rn*>) (A), *E93*-depleted (*rn>E93$^{RNAi}$*) (B), and *E93*-depleted with simultaneous overexpression of the inhibitor of caspases *P35* (*rn>E93$^{RNAi}$; P35*) (C) of 24h APF pupal wings stained with DAPI to visualize nuclei, and anti-Dcp-1 to visualize caspase activity. (D-F) Representative images of adult wings of the indicated genotypes. (G) Measurement of adult wings area of the indicated genotypes. Error bars indicate the SEM (n = 7-12). Asterisks indicate differences statistically significant at ***$p \leq 0.001$ (*t*-test). Pupal wing contours are labelled by white lines. Scale bars: 100 μm.

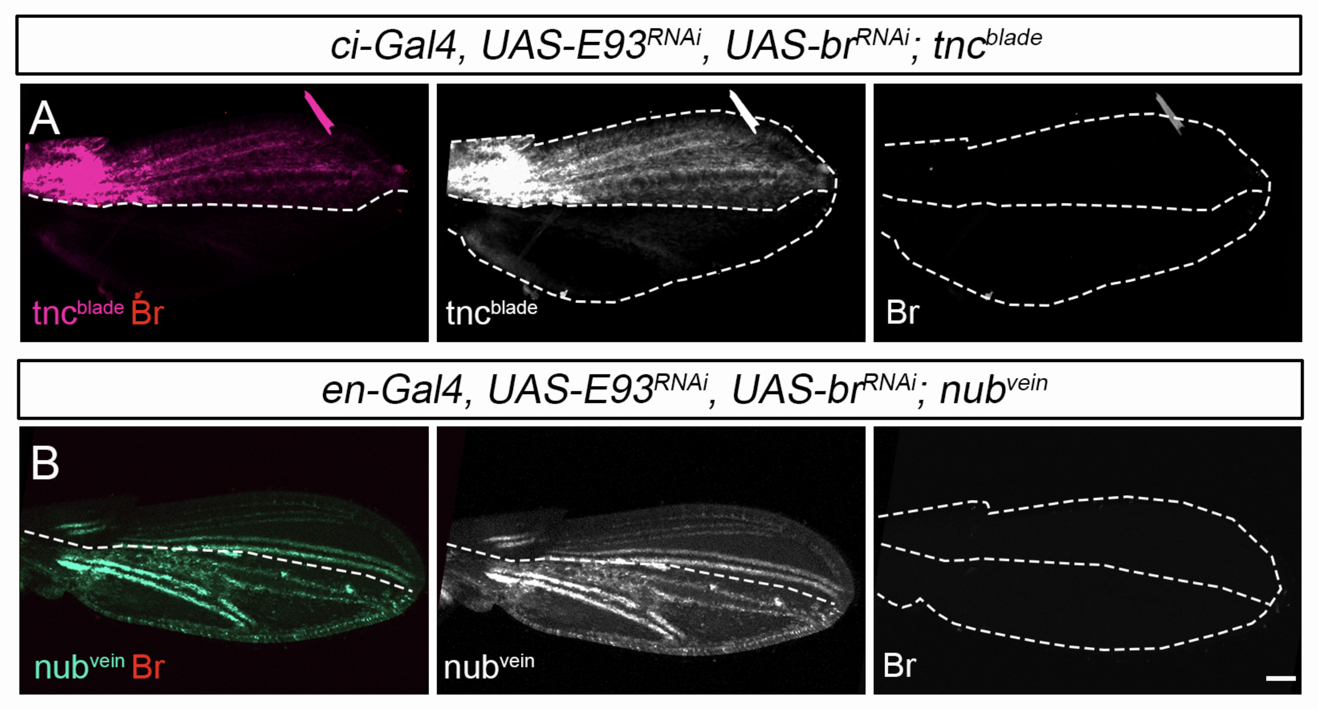


**Figure S3. The activation of E93-dependent pupal-specific enhancers is mediated by *br* repression.** Confocal images of *tnc*$^{blade}$ (A) and *nub*$^{vein}$ (B) enhancer activities in pupal wings simultaneously expressing *E93*$^{RNAi}$ and *br*$^{RNAi}$ under the control of *ci-Gal4* or *en-Gal4*, respectively. Wings were stained with anti-Br to visualize the absence of expression of Br in the *E93/Br-depleted* domains. Dotted lines indicate the contours of the pupal wings and the boundary of the corresponding driver expression domain. First panels are overlay images of the corresponding panels on the right. Scale bar: 50 μm.

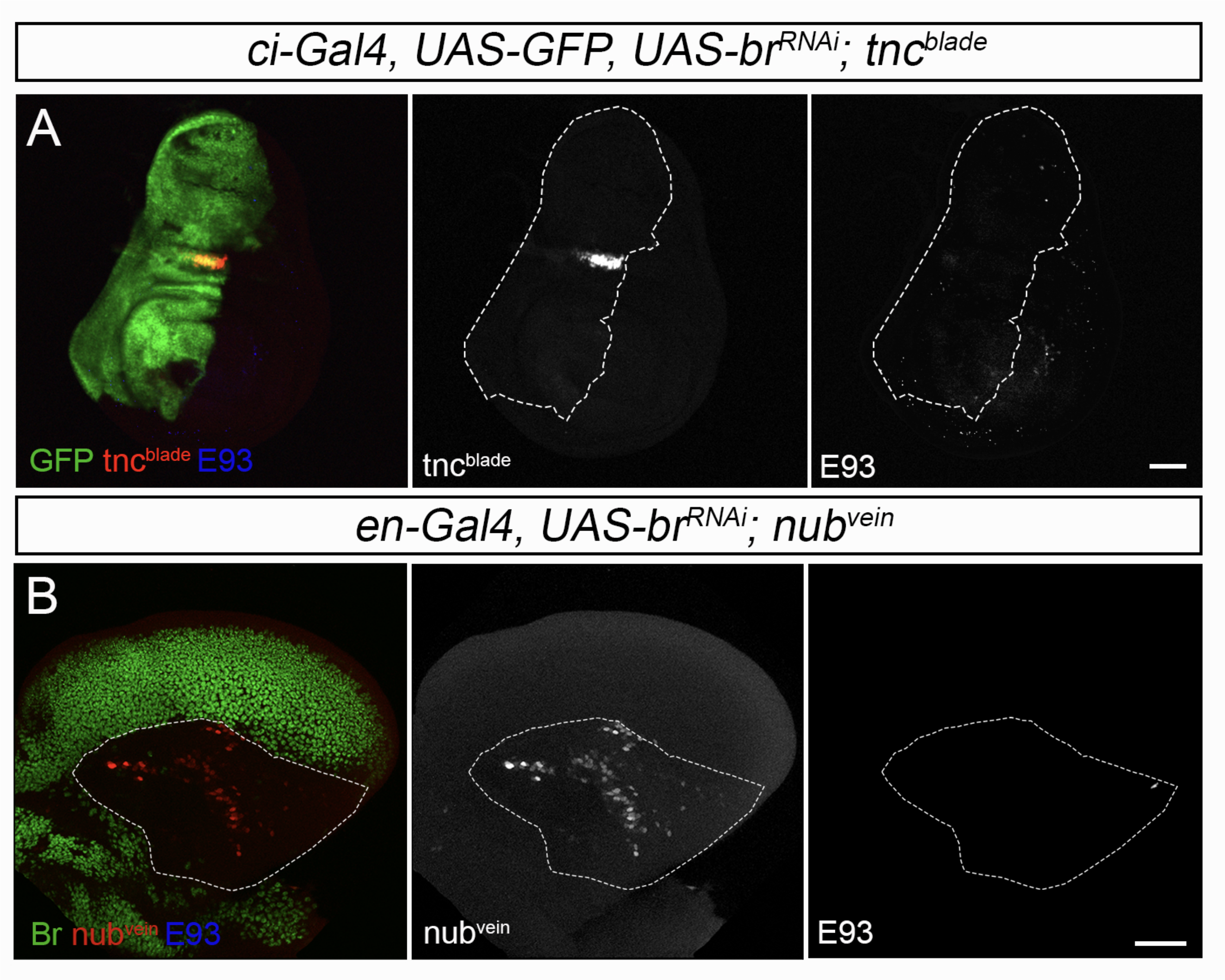


**Figure S4. Br suppresses the activity of pupal-enhancers in larval wing discs and prepupal wings.** (A) Confocal images of *tnc*$^{blade}$ enhancer activity, and immunostaining of E93 in wandering L3 wing discs expressing *br*$^{RNAi}$ under the control of *ci-Gal4*. (B) Confocal images of *nub*$^{vein}$ enhancer activity and immunostaining of E93 in 9h prepupal wings expressing *br*$^{RNAi}$ under the control of *en-Gal4*. Dotted lines indicate the boundary of the corresponding driver expression domain. First panels are overlay images of the corresponding panels on the right. Scale bars: 50 μm.
